# Analysing disturbance response of cell-to-cell communication systems: a case study for activator-repressor-diffuser motif

**DOI:** 10.1101/304436

**Authors:** Taishi Kotsuka, Yutaka Hori

## Abstract

In biomolecular communication networks, bacterial cells communicate with each other using a cell-to-cell communication mechanism mediated by diffusible signaling molecules. The dynamics of molecular concentrations in such systems are approximately modeled by reaction-diffusion equations. In this paper, we analyse the ability of cell-to-cell communication systems to attenuate impulsive disturbances with various spatial frequency profiles by computing the integrated squared concentration of molecules. In particular, we perform in-depth study of disturbance responses for an activator-repressor-diffuser biocircuit in the spatial frequency domain to characterize its spatial frequency gain.

## 1 Introduction

Bacterial cells implement a communication mechanism by secreting and sensing diffusible signaling molecules called autoinducer [1, 2]. The cell-to-cell communication mechanism, which is called quorum sensing allows bacteria to regulate gene expression at a population level and achieve coordinated behaviors. Examples of coordinated behaviors range from the formation of biofilm and the production of antibiotics to the regulation of bioluminescence [3]. Since these phenomena often cause significant impact in our daily life both in positive and negative ways [2], it is important to understand the underlying mechanism of the cell-to-cell communication.

In synthetic biology, cell-to-cell communication has been actively utilized as a mechanism to implement feedback loops between cells. Specifically, synthetic biologists engineered biomolecular communication networks that enable multicellular computing [4, 5], synchronization [6, 7, 8] and pattern formation [9, 10, 11, 12]. The importance of robust cell-to-cell communication has been particularly increasing in recent years to implement many steps of reactions that cannot be built in a single cell due to the short of cellular resource.

To enable robust engineering of multicellular biocircuits and better understand the underlying mechanism, this paper presents an approach to quantitatively analysing the effect of disturbance, or noise, on cell-to-cell communication based on a reaction-diffusion model of molecular communication system. Specifically, we compute the total pseudo-energy of the disturbance that propagates in the cell-to-cell communication system based on the *H*_2_ norm computation method for linear time-invariant (LTI) system [13]. Our analysis allows facile characterization of the bandwidth of the noise frequency that a given cell-to-cell communication attenuates/amplifies. Thus, it becomes a useful guidance for the robust design of biomolecular communication networks. We apply the proposed approach to the activator-repressor-diffuser biomolecular circuit [14] to find a circuit configuration that filters global disturbance to the system and discuss a feasible implementation of such biomolecular circuit.

The organization of this paper is as follows. In Section 2, we introduce a model of biomolecular communication network. In Section 3, we present an algebraic equation to compute the spatial frequency response. Section 4 analyses the disturbance response of a synthetic biomolecular communication network, and discusses the analysis result in the context of synthetic biology. Finally, Section 5 concludes this paper.

## 2 Model of Biomolecular Communication Networks

We consider molecular communication networks consisting of *n* molecular species in one dimensional space Ω := [0, *L*] for notational simplicity. We assume that a large number of cells communicate as illustrated in Figure 1, and cells produce transmitter molecules that diffuse in the spatial domain. Let *x_i_*(*t, ξ*) denote the concentration of the *i-*th molecule at position *ξ* ∈ Ω and at time *t*, and define *x*(*t, ξ*) := [*x*_1_(*t, ξ*),*x*_2_(*t, ξ*), …, *x_n_*(*t, ξ*)]^*T*^. The dynamics of the concentrations of molecular species are formulated by

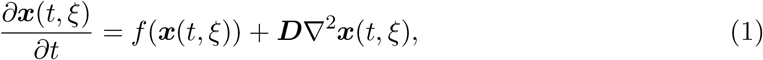

where *f* (·) is a function representing the rate of production and degradation of the molecules [14]. *D* is a diagonal matrix of diffusion constants. Note that, in practice, biomolecular systems use a single diffusible molecules, thus *D* has a single non-zero diagonal entry. In what follows, we assume the Neuman boundary condition as follows.

**Figure 1:**
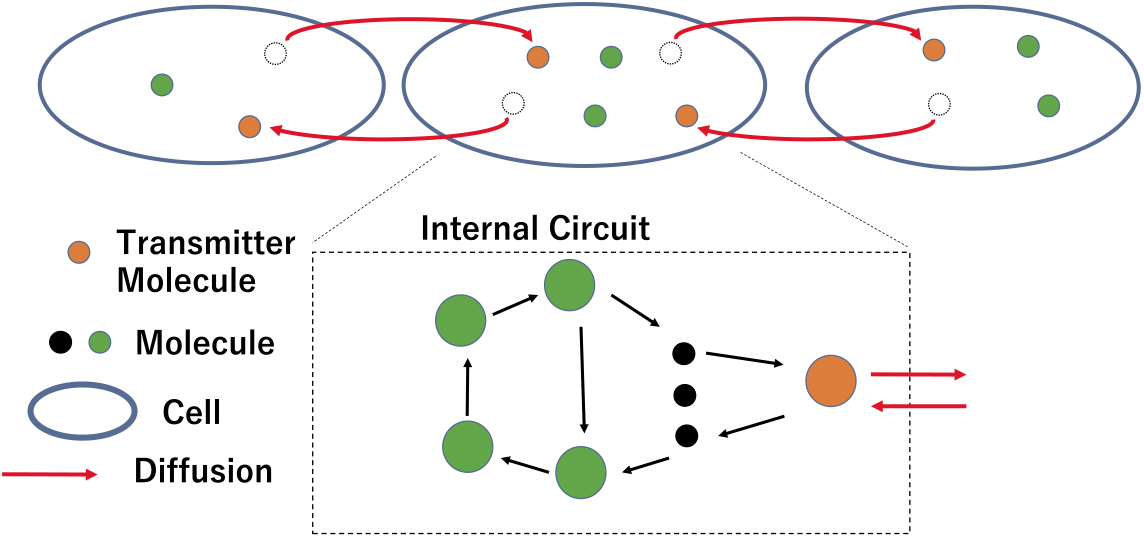
Molecular communication network. The chemical concentrations inside cells are approximately modeled by the continuous gradient of concentrations *x*(*t, ξ*).

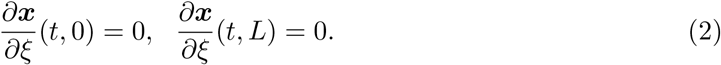

Let 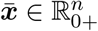 denote a spatially homogeneous equilibrium state of a single cell. The local dynamics around 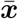 are modeled by the following linearized model of (1).

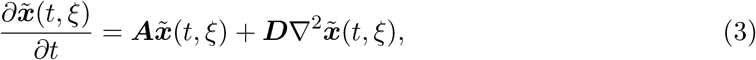

where 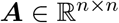 is the Jacobian of *f* (·) at 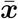, and 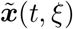 is defined by 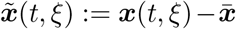. Note that the system (3) is an infinite dimensional linear time-invariant system.

## 3 Disturbance response analysis in spatial frequency domain

In this section, we first define a norm of disturbance response of molecular communication network. We then show that the reaction-diffusion system (1) can be decomposed into subsystems describing the dynamics of spatial modes. This decomposition facilitates the spatial frequency domain characterization of the disturbance response.

We consider to analyse disturbance response of molecular communication networks against impulsive perturbation. More specifically, we consider impulse perturbation

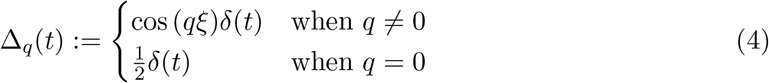

with Dirac delta function *δ*(*t*) and compute the *L*^2^ norm

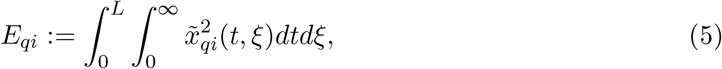

where 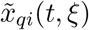 is the *i*-th molecular concentration when the profile of the perturbation is given by, Δ*_q_*(*t*), which is a sinusoid with spatial frequency *q*.

It should be noted that *E_qi_* is the integral of the squared concentration, which can be considered as a total pseudo-energy of *x_i_*(*t, ξ*) when the cell-to-cell communication system undergoes spatially periodic perturbation Δ*_q_*(*t*). Since

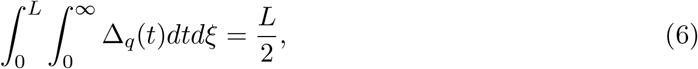

2*E_qi_*/*L* gives the gain of the energy for the impulsive perturbation with spatial frequency *q*. The gain provides a quantitative measure of disturbance attenuation/amplification by a molecular communication network.

In what follows, we show that the infinite dimensional system (3) can be decomposed into finite dimensional subsystems that account for the dynamics of individual spatial modes. Using the decomposed system, we characterize the integrated squared concentration *E_qi_*.

Fourier series expansion of the molecular concentrations 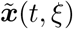 can be expressed as

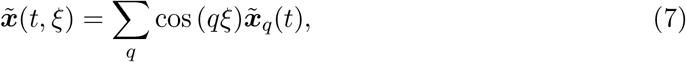

where *q* = *kπ/L* (*k* = … -1, 0, 1, …). We define the harmonic component of the spatial frequency *q* as 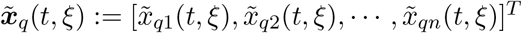, that is,

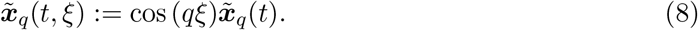

It should be noticed that the *i*-th entry of 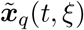 is 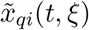 defined in Eq. (5). Substituting Eq. (7) into Eq. (3) and extracting the harmonic component 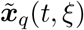, the dynamics of molecular concentration 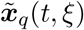 are obtained as

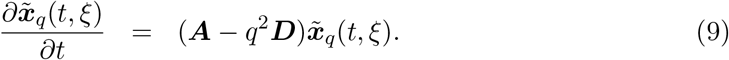

Eq. (9) implies that the temporal dynamics of each spatial harmonic component are modeled by a finite order ordinary differential equation (ODE). Thus, we can compute the squared concentration *E_qi_* by analysing the solution to the ODE (9).

### Theorem 1.

Consider the reaction-diffusion system (3) and assume ***A*** — *q^2^**D*** is Hurwitz. The *i*-th diagonal entry of *P_q_* that satisfies

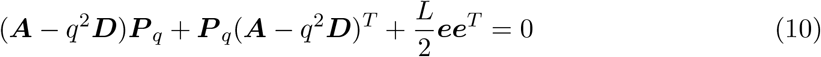

is *E_qi_*, where we define 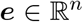 as a vector that consists of 0 and 1. The *i*th entry of *e* is 1 when the impulse perturbation is input into the *i*-th molecule.

Theorem 1 is based on the well-known Lyapunov equation for *H*_2_ norm computation in control engineering [13]. Since Eq. (10) is linear in the variable *P_q_*, the squared integral *E_qi_* defined in (5) can be efficiently computed without simulating the LTI system (9).

## 4 Disturbance Response of Activator-Repressor-Diffuser System

In this section, we use Theorem 1 to analyse the disturbance response of a synthetic biomolecular communication network called activator-repressor-diffuser system proposed in [14]. We then interpret the analysis result in the context of synthetic biology.

### 4.1 Spatial Frequency Response Analysis

We consider activator-repressor-diffuser system [14] in Figure 2, where the three molecules, activator, repressor and diffuser, form a reaction network in each cell, and cells communicate with each other by exchanging the diffuser molecule. The dynamics of the system are modeled by

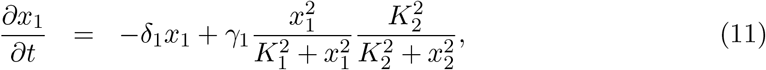

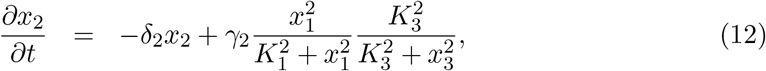

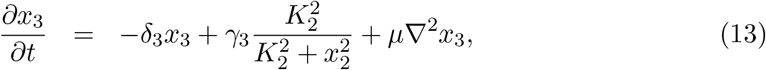

where *x*_1_, *x*_2_ and *x*_3_ are the concentrations of activator, repressor and diffuser, respectively. *δ_i_* and *γ_i_* are the degradation rate and the production rate of the *i*-th molecule, respectively, and *K_i_* is the Michaelis Menten constant, and *μ* is the diffusion constant. We assume that the impulse perturbation is input into all molecules in the cells. *e* is thus obtained as *e* := [1,1,1]^*T*^. To analyse the spatial frequency response of the activator-repressor-diffuser system to the impulse perturbation, we numerically solve the Lyapunov equation (10) for different spatial frequency *q* using the parameter sets shown in Table 1, where the degradation rate *δ*_1_ is varied between 0.001 min^−1^ and 0.05 min^−1^. These parameter values are consistent with widely used parameters for numerical simulations of biomolecular reactions in E. coli [14]. We set the diffusion coefficient as *μ* = 8.6 × 10^−8^ mm^2^min^−1^, which is consistent with literature values [15]. The length of the reaction domain is set *L* = 3.0 mm.

**Figure 2:**
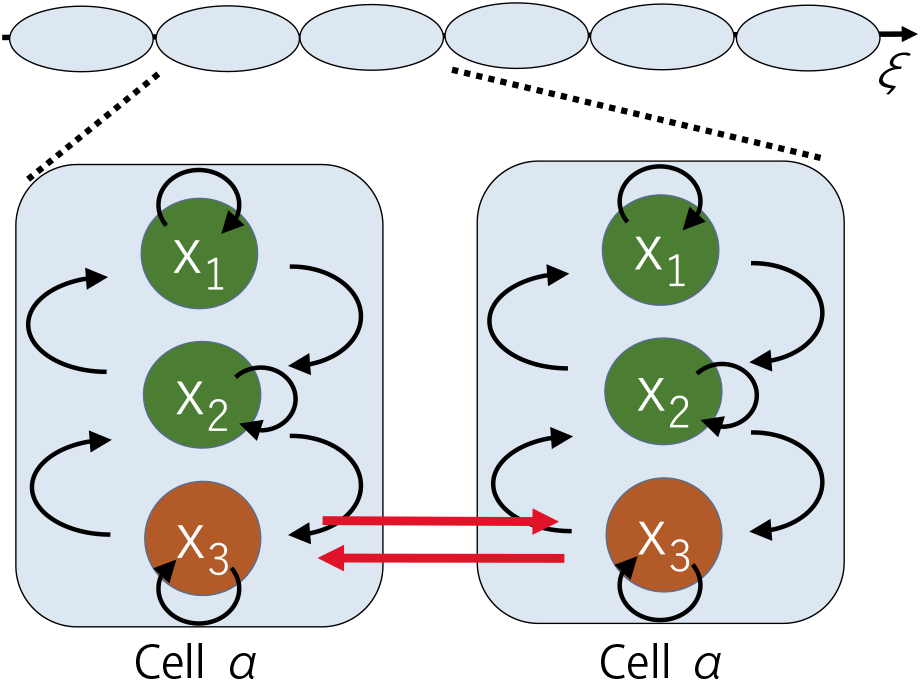
Molecular communication network of activator-repressor-diffuser system.

**Table 1:**
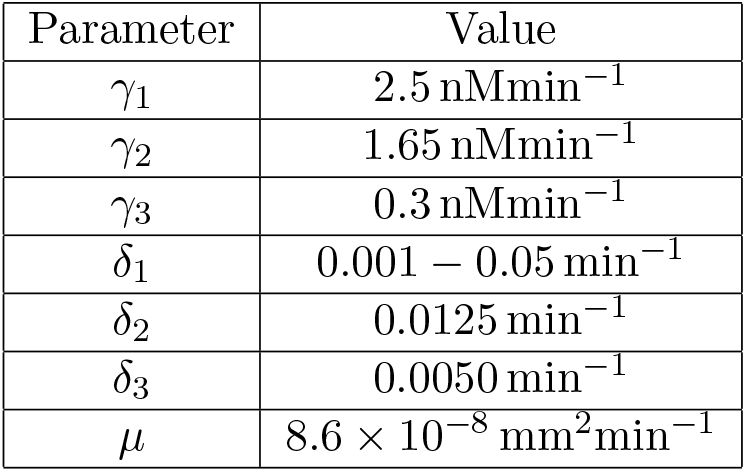
The parameter sets for the analysis of the model Figure 2

Figure 3 illustrates the computation result of the integrated squared concentration *E_q1_*

**Figure 3:**
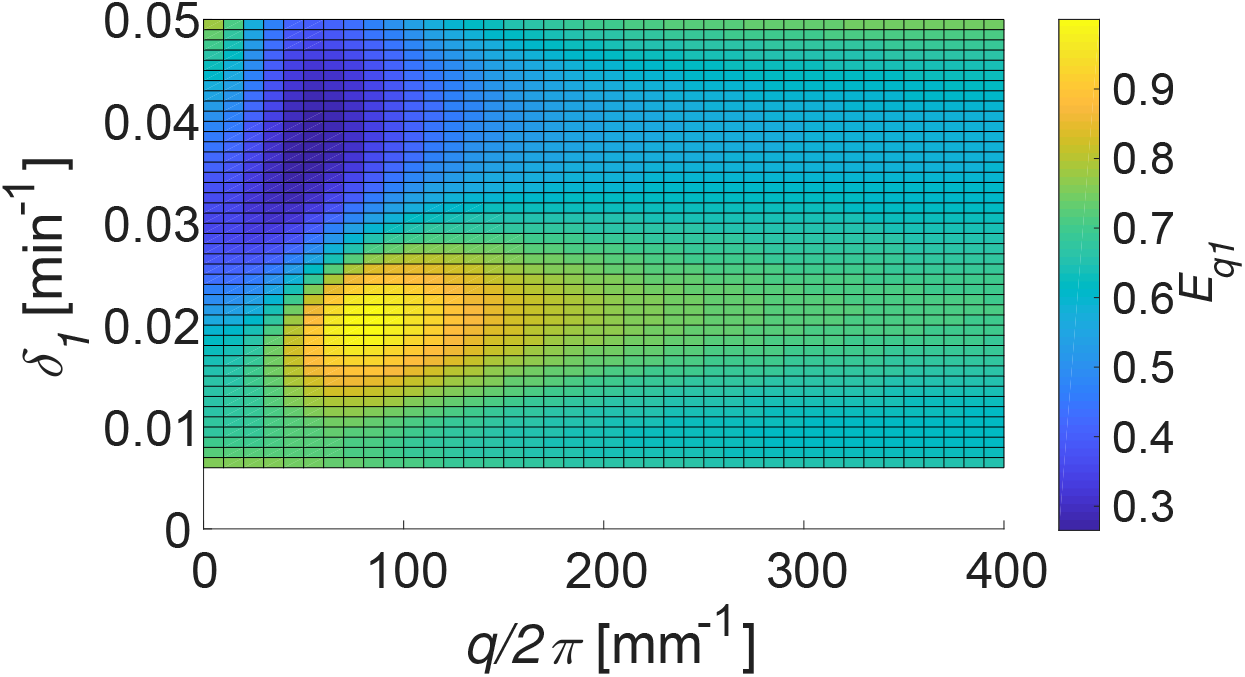
Integrated squared concentration of *x_q1_* for different degradation rate *δ*_1_ and *q/2π*. The colormap is normalized with the maximum value of *E_q1_* set to 1.

The figure shows two types of response characteristics for *q/2π* ≤ 150 mm^−1^. Specifically, the value of *E_qi_* is larger between *q/2π* ≤ 150 mm^−1^ than the other frequency range when *δ*_1_ = 0.02 min^−1^, while it is smaller when *δ*_1_ = 0.04 min^−1^. This implies that the disturbance response of the activator-repressor-diffuser system shows qualitatively different behavior, *i.e.*, amplification or attenuation, depending on the degradation rate of the activator molecule.

### 4.2 Interpretation and discussion

In what follows, we perform in-depth study of the computation result in Figure 3 to elucidate the effect of cell-to-cell communication on the disturbance response. Based on this analysis, we discuss feasible design of biomolecular communication networks that can attenuate the effect of disturbance.

Figure 4 illustrates the integrated squared concentration *E_q1_* for *δ*_1_ = 0.02 min^−1^ and *δ*_1_ = 0.04min^−1^. The red line in Figure 4 is the value of *E_q1_* when there is no cell-to-cell communication, which we can calculate from Eq. (10) with *D* = 0. Figure 4 clearly illustrates that cell-to-cell communication affects the disturbance response of molecular communication network systems.

**Figure 4:**
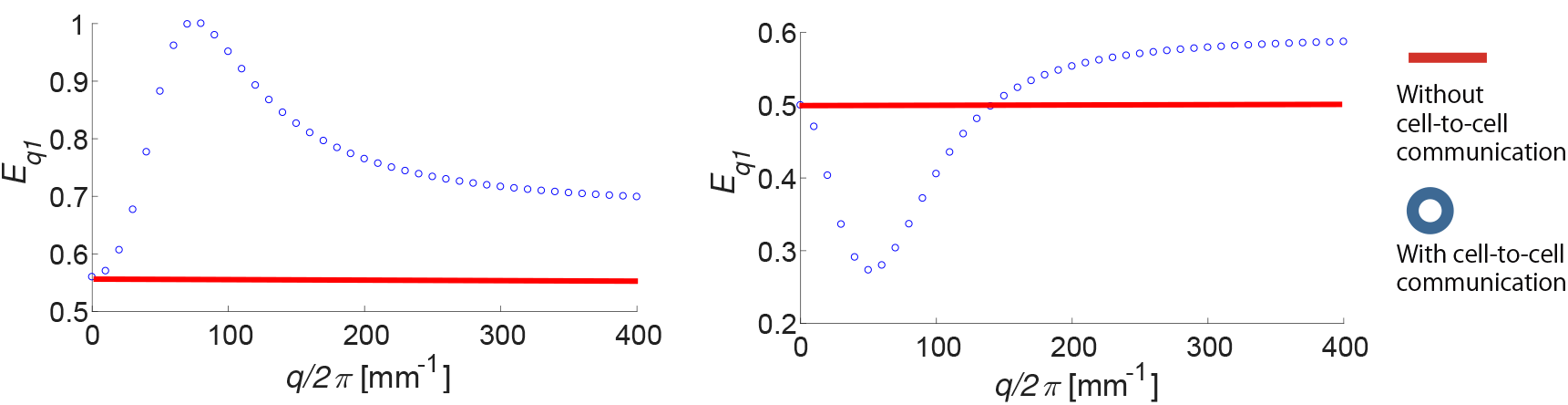
Integrated squared concentration (A) for the degradation rate *δ*_1_ = 0.02. (B) for the degradation rate *δ*_1_ = 0.04.

Recall that *E_q1_* is the index showing how much the perturbation Δ(*t*) excites *x_q_*(*t, ξ*). Figure 4 (A) shows that cell-to-cell communication amplifies the disturbance of *x*_1_ when *δ*_1_ = 0.02min^−1^, while Figure 4 (B) shows that the communication attenuates the disturbance when the bandwidth of the disturbance is *q/2π* ≤ 150mm^−1^. To interpret this result, we compare the bandwidth with the size of E. coli cells, which is around 1.0 *μ*m. The reciprocal of *q/2π* ≤ 150mm^−1^ leads to the spatial wave length of the disturbance as *q/2π* ≥ 6.66 *μ*m, which is about the size of seven cells. This means that the cell-to-cell communication of the activator-repressor-diffuser system attenuates the disturbance that globally affects the gene expression at the population level, but it amplifies single-cell-level disturbance.

The parameter *δ*_1_ accounts for the lifetime of the protein, which is determined by both degradation and the division of a cell. From a design point of view, tuning of the rate parameter *δ*_1_ is possible by adding a degradation tag to the protein for active degradation, or changing the growth condition such as cell culture medium to moderate the division rate. The generation time of E. coli is normally around 20 minutes in LB medium, but it can be controlled from 20 to 80 minutes by changing the culture medium [16, 17, 18]. The degradation rate *δ*_1_ = 0.04min^−1^ is about 20 minutes of half-life. This implies that *δ*_1_ = 0.04min^−1^ is a feasible parameter that we can tune by growing the cells in LB medium.

## 5 Conclusion

We have analysed the disturbance response of cell-to-cell communication systems based on the reaction-diffusion model with a single diffuser. We have first shown that the infinite dimensional system can be decomposed into finite dimensional subsystems that account for the dynamics of spatial modes. We have then introduced an approach to analysing the pseudo-energy that propagates in the system when a unit impulse perturbation is applied. Applying the proposed approach to the activator-repressor-diffuser system, we have characterized the reaction parameters with which a broad spatial bandwidth of noise can be attenuated. Finally, we have verified based on the existing literatures that such parameter values are feasible in wet-lab implementation.

## Acknowledgments

This work was supported in part by the Okawa Foundation Research Grant under grant number 16-10.

## References

[1] C. M. Waters and B. L. Bassler, “Quorum sensing: cell-to-cell communication in bacteria,” Annual Review of Cell and Cevelopmental Biology, vol. 21, pp. 319–346, 2005.

[2] S. J. Hagen, The physical basis of bacterial quorum communication. Springer, 2014.

[3] V. C. Thomas, L. R. Thurlow, D. Boyle, and L. E. Hancock, “Regulation of autolysis-dependent extracellular DNA release by enterococcus by enterococcus faecalis extracellular proteases influences biofilm development,” Journal of Bacteriology, vol. 190, no. 16, pp. 5690–5698, 2008.

[4] T. S. Moon, C. Lou, A. Tamsir, B. C. Stanton, and C. A. Voigt, “Genetic programs constructed from layered logic gates in single cells,” Nature, vol. 491, no. 7423, p. 249,2012.

[5] A. Tamsir, J. J. Tabor, and C. A. Voigt, “Robust multicellular computing using genetically encoded NOR gates and chemical ‘wires’,” Nature, vol. 469, no. 7329, p. 212,2011.

[6] T. Danino, O. Mondragón-Palomino, L. Tsimring, and J. Hasty, “A synchronized quorum of genetic clocks,” Nature, vol. 463, no. 7279, p. 326,2010.

[7] M. O. Din, T. Danino, A. Prindle, M. Skalak, J. Selimkhanov, K. Allen, E. Julio, E. Atolia, L. S. Tsimring, S. N. Bhatia, et al., “Synchronized cycles of bacterial lysis for in vivo delivery,” Nature, vol. 536, no. 7614, p. 81,2016.

[8] B. Leo, M. Mather, and H. Jeff, “Synchronized DNA cycling across a bacterial population,” Nature Genetics, vol. 49, no. 8, pp. 1282–1285, 2017.

[9] S. R. Scott, M. O. Din, P. Bittihn, L. Xiong, L. S. Tsimring, and J. Hasty, “A stabilized microbial ecosystem of self-limiting bacteria using synthetic quorum-regulated lysis,” Nature microbiology, vol. 2, no. 8, p. 17083, 2017.

[10] L. You, R. S. Cox III, R. Weiss, and F. H. Arnold, “Programmed population control by cell-cell communication and regulated killing,” Nature, vol. 428, no. 6985, p. 868,2004.

[11] S. Basu, Y. Gerchman, C. H. Collins, F. H. Arnold, and R. Weiss, “A synthetic multicellular system for programmed pattern formation,” Nature, vol. 434, no. 7037, p. 1130, 2005.

[12] C. Liu, X. Fu, L. Liu, X. Ren, C. K. Chau, S. Li, L. Xiang, H. Zeng, G. Chen, L.-H. Tang, et al., “Sequential establishment of stripe patterns in an expanding cell population,” Science, vol. 334, no. 6053, pp. 238–241, 2011.

[13] K. Zhou, J. C. Doyle, and K. Glover, Robust and optimal control. Prentice hall New Jersey, 1996.

[14] Y. Hori, H. Miyazako, S. Kumagai, and S. Hara, “Coordinated spatial pattern formation in biomolecular communication networks,” IEEE Transactions on Molecular Biological and Multi-Scale Communications, vol. 6, no. 2, pp. 111–121, 2015.

[15] T. J. Lardner, “The measurement of cell membrane diffusion coefficients,” Jounal of Biomechanics, vol. 10, pp. 167–170, 1977.

[16] G. Sezonov, D. Joseleau-Petit, and R. D’Ari, “Escherichia coli physiology in Luria-Bertani broth,” Journal of Bacteriology, vol. 189, no. 23, pp. 8746–8749, 2007.

[17] N. Peekhaus and T. Conway, “What’s for dinner?: Entner-Doudoroff metabolism in Escherichia coli,” Journal of Bacteriology, vol. 180, no. 14, pp. 3495–3502, 1998.

[18] H. Tao, C. Bausch, C. Richmond, F. R. Blattner, and T. Conway, “Functional genomics: expression analysis of Escherichia coli growing on minimal and rich media,” Journal of Bacteriology, vol. 181, no. 20, pp. 6425–6440, 1999.

